# FAM21 is critical for TLR2-CLEC4E-mediated dendritic cell function against *Candida albicans*

**DOI:** 10.1101/2022.02.16.480635

**Authors:** Rakesh Kulkarni, Siti Khadijah Kasani, Ching-Yen Tsai, Shu-Yun Tung, Kun-Hai Yeh, Chen-Hsin Albert Yu, Wen Chang

## Abstract

FAM21 (family with sequence similarity 21) is a component of the Wiskott–Aldrich syndrome protein and SCAR homologue (WASH) protein complex that mediates actin polymerization at endosomal membranes to facilitate sorting of cargo-containing vesicles out of endosomes. To study the function of FAM21 *in vivo*, we generated conditional knockout (cKO) mice in the C57BL/6 background in which FAM21 was specifically knocked out of CD11c-positive dendritic cells. Bone marrow-derived dendritic cells (BMDC) from those mice displayed enlarged early endosomes, and altered cell migration and morphology relative to wildtype (WT) cells. FAM21-cKO cells were less competent in phagocytosis and antigen processing *in vitro*, though antigen presentation was not affected. More importantly, we identified the TLR2/CLEC4E signaling pathway as being downregulated in FAM21-cKO BMDCs when challenged with its specific ligand *Candida albicans*. Moreover, FAM21-cKO mice were more susceptible to *C. albicans* infection than WT mice. Reconstitution of WT BMDCs in FAM21-cKO mice rescued them from lethal *C. albicans* infection. Thus, our study highlights the importance of FAM21 in a host immune response against a significant pathogen.

## Introduction

Endocytosis is defined as the uptake of components on the plasma membrane and in extracellular fluids into the cytoplasm (Kaksonen & Roux, 2018). Several types of endocytosis have been identified, including clathrin-mediated endocytosis (CME), caveolae, macropinocytosis, phagocytosis, and clathrin-independent pathways, which differ depending on the size and type of cargo that is endocytosed (Doherty & McMahon, 2009). Endocytosis mediates important biological processes such as nutrient uptake, cell signalling, antigen presentation, cell adhesion, migration and mitosis (Doherty & McMahon, 2009). Pathogens such as bacteria and viruses exploit the cellular endocytic machinery to enter host cells (Bonazzi & Cossart, 2006; Brass *et al*, 2008; Cossart & Helenius, 2014). Several studies have highlighted the role of the actin cytoskeleton in regulating endocytosis such as via recruitment of adaptor proteins, concentration of cargo, stabilization of endosomal domains, and formation of phagocytic cups (Galletta & Cooper, 2009; Kumari *et al*, 2010; Simonetti & Cullen, 2019). Depending on their function, cargoes present in endosomes are directed towards their respective destinations (Deng *et al*, 2015; Mooren *et al*, 2012). Mechanisms of endocytosis and cellular trafficking are orchestrated by multi-protein assemblies that include retromer and retriever, sorting nexins, and the ARP2/3 activating WASH complex (Fokin & Gautreau, 2021; Seaman, 2012; Simonetti & Cullen, 2019). Our previous studies on HeLa cells have shown that vaccinia mature virus (VACV) is endocytosed into host cells and that a novel protein, FAM21- also known as vaccinia virus penetration factor (VPEF), is required for virus penetration (Hsiao *et al*, 2015). Knockdown (KD) of FAM21 expression blocked intracellular transport of dextran and VACV penetration into cells but did not inhibit their binding to cells, suggesting that FAM21 mediates the VACV endocytic process in HeLa cells (Huang *et al*, 2008).

FAM21 is a large 1334-amino acid (aa) protein, containing a globular ‘head’ domain (∼200 aa) at its N-terminus and a long (∼1100 aa) C-terminal ‘tail’ that contains a binding site for the actin-capping protein CapZ (Gomez & Billadeau, 2009; Jia *et al*, 2012). FAM21 is a component of the WASH protein complex that also comprises WASH1, Strumpellin, Strumpellin and WASH interacting protein (SWIP), and Coiled-coil domain-containing protein 53 (CCDC53) (Harbour *et al*, 2012; Jia *et al*., 2012). The WASH protein complex interacts with endosomes through the binding of its constituent FAM21 to the vacuolar protein sorting retromer complex (consisting of VPS26, VPS29 and VPS35) anchored on endosomal membranes (Harbour *et al*., 2012). The WASH protein complex then recruits actin-related protein 2/3 (ARP2/3) to mediate actin polymerization at endosomal membranes so that cargo-containing vesicles can be budded out of endosomes (Gomez & Billadeau, 2009). FAM21 is also required for trafficking of receptor proteins essential for nutrient uptake, such as the glucose transporter GLUT1 (Lee *et al*, 2016) and the copper transporter Copper metabolism MURR1 domain-containing 1 (COMMD1) (Phillips-Krawczak *et al*, 2015). When impaired, disruption of these receptor proteins results in deficiencies of essential cell nutrients.

Although most FAM21 functions in endosomal sorting and trafficking are dependent on the WASH protein complex(Gomez & Billadeau, 2009), previous studies have shown that FAM21 also exerts WASH-independent functions (Deng *et al*., 2015; Lee *et al*., 2016). Studies have shown that FAM21, but not the WASH complex, could enhance nuclear factor kappa B (NF-κB) transcriptional activation (Deng *et al*., 2015; Lee *et al*., 2016). Another study by Lee et al. (2016) showed that FAM21 regulates PI4KB levels at Golgi independently of WASH complex (Lee *et al*., 2016). In mammalian cells, but not in *Dictyostelium* (Park *et al*, 2013), formation of intact WASH complex dictates the stability of each of its subunits (Derivery *et al*, 2009; Jia *et al*., 2012), and it has been shown that an E3 RING ubiquitin ligase, MAGE-L2-TRIM27, targets WASH protein for degradation (Hao *et al*, 2013). K63 ubiquitination of WASH at residue K220 is facilitated by MAGE-L2-TRIM27, which results in F-actin nucleation and retromer-dependent transport (Hao *et al*., 2013).

The trafficking defects derived from mutations or loss of functional subunits in the WASH complex results in a wide range of human diseases. The WASH complex has been demonstrated as important for cell invasion by *Salmonella* (Hanisch *et al*, 2010; Unsworth *et al*, 2004). Mutations in Strumpellin cause autosomal dominant hereditary spastic paraplegia, a neurodegenerative disorder of upper neurons (Valdmanis *et al*, 2007). Homozygous P1019R mutation in SWIP causes autosomal-recessive mental retardation 43 (MRT-43) (Ropers *et al*, 2011). A point mutation (D620N) within the VPS35 subunit of the vacuolar protein sorting retromer complex, with which the WASH complex directly interacts, is associated with Parkinson’s disease (PD) and is also responsible for disrupting cargo sorting (Follett *et al*, 2014). Previous studies have also shown that KO of WASH, Strumpellin and the retromer protein subunit VPS26a all individually result in embryonic lethality (Muhammad *et al*, 2008; Piotrowski *et al*, 2013; Tyrrell *et al*, 2016), though VPS26b-KO mice survived despite displaying sortillin trafficking defects (Kim *et al*, 2010). Genetic disruption arising from SWIP^P1019R^ mutation resulted in significant and progressive motor deficits in mice that were similar to movement deficits observed in humans (Courtland *et al*, 2021). Despite being a component of the WASH protein complex, whether FAM21 mutations are associated with any known human diseases remains unknown, so our investigation of the function of FAM21 *in vivo* is critical to understanding if FAM21 plays a role in disease prevention.

## Results

### FAM21 cKO BMDCs exhibit enlarged early endosomes, as well as reduced cell spread and migration

To investigate the role of FAM21 *in vivo*, we generated a constituitive FAM21 KO mouse line (Figure EV1A), but no FAM21^-/-^ littermates were obtained. FAM21 deficiency caused embryonic lethality as early as E7.5 (Figure EV1B), evidencing an essential role of FAM21 protein in embryonic development. Northern blot analyses revealed that FAM21 is widely expressed in multiple organs, including brain, intestine, kidney, liver, lungs, lymph nodes and thymus (Figure EV1C). FAM21 is also expressed in BMDCs (Figure EV1C), allowing us to address its functions in immune responses by generating mice with a floxed *FAM21* gene (Figure EV1D). We then crossed the FAM21^f/+^ mice with E2a-Cre mice (Figure EV1E) to obtain heterozygous FAM21^+/-^ mice that could be used to knock out the *FAM21* gene in CD11c-positive dendritic cells in order to study the roles of FAM21 in BMDCs *in vitro* (Figure EV1E). However, that strategy generated KO mice unsuited to *in vivo* experiments, so we bred Fam21^f/f^ mice with CD11c-Cre mice to specifically knock out the *FAM21* gene in CD11c-positive dendritic cells to address the biological roles of FAM21 *in vivo* (in Figure EV1F). All mice generated according to these two KO strategies developed normally with no obvious phenotypic differences (data not shown).

Bone marrow cells extracted from femur and tibia of WT and the cKO mice described above were cultured with GM-CSF to induce dendritic cell differentiation *in vitro*. At day 8, the cells were analyzed using anti-CD11b and anti-CD11c antibodies, which revealed similar levels (∼85%) of CD11c^+^CD11b^+^ BMDCs in both WT and cKO mice. Less than 10% of the BMDC cultures expressed high levels of CD86 and MHC class II activation markers, indicating that the majority were immature BMDCs (Figure 1A). Both RT-PCR (Figure 1B) and immunoblot analysis (Figure 1C) confirmed that FAM21 expression was significantly reduced in FAM21-cKO BMDCs. Although FAM21 protein displayed three forms of different molecular weight, perhaps derived from alternative splicing (Figure EV2), all three forms were significantly reduced in the cKO BMDCs (Figure 1C).

**Figure 1:**
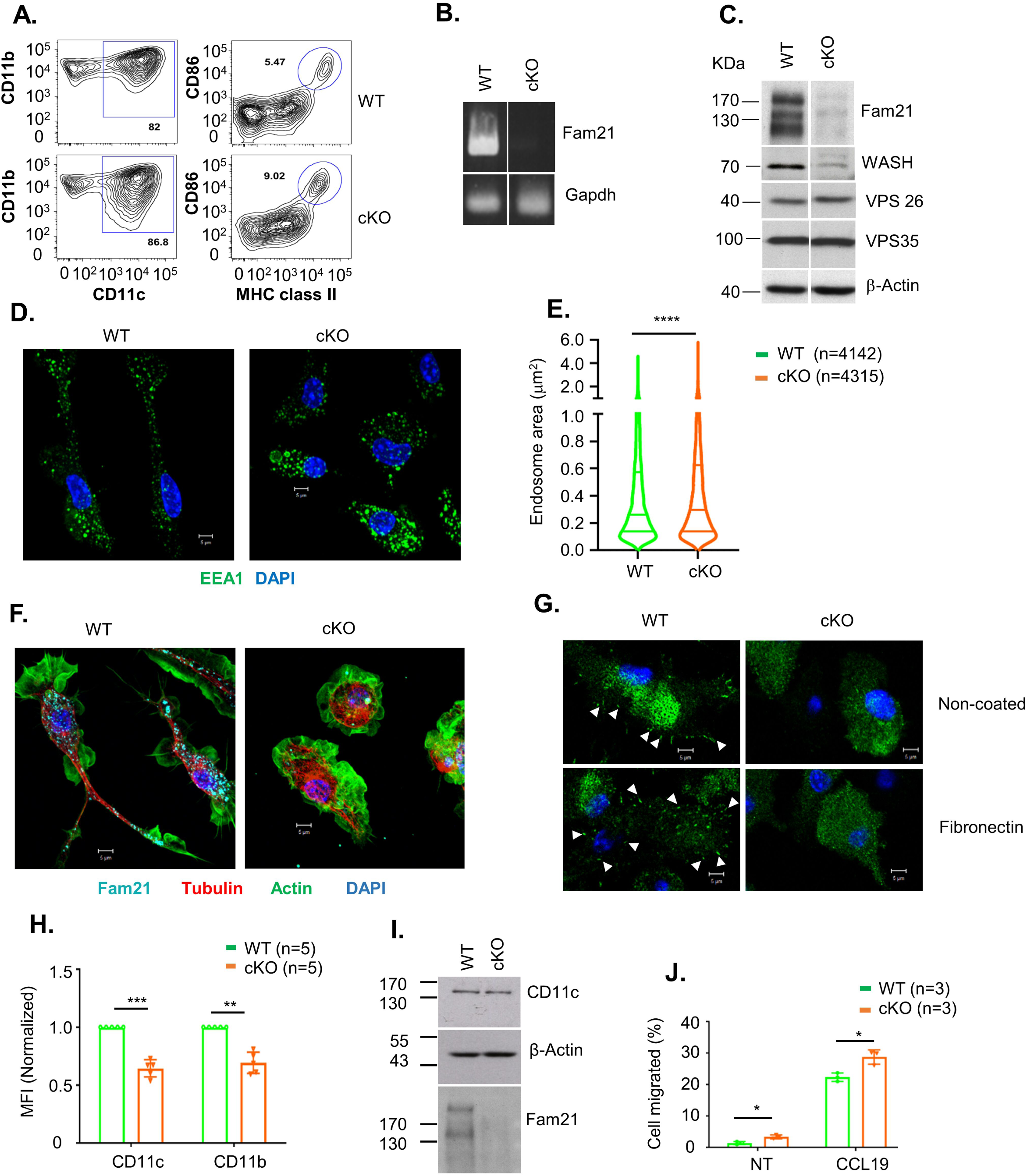
FAM21-cKO BMDCs present enlarged early endosomes and exhibit reduced cell spreading ability. (A) Flow cytometry of BMDCs isolated at day 8 post isolation from WT and FAM21-cKO mice using anti-CD11b, anti-CD11c, anti-CD86 and anti-MHC class II antibodies. (B) RT-PCR of FAM21 transcript in WT and cKO BMDCs. GAPDH was used as a control. (C) Immunoblot of FAM21 in BMDCs cultured from WT and FAM21-cKO mice. WASH and the retromer complex proteins VPS26 and VPS35 are also shown. β-Actin was used as a control. (D) Immunofluorescence staining of early endosomes (green), and nuclei (DAPI, blue) in WT and cKO BMDCs. (E) Quantification of endosome size distributions in WT and cKO BMDCs using violin plots. n = total number of endosomes measured for each genotype. ****p<0.0001. (F) Confocal images of WT and cKO BMDCs stained with anti- FAM21 (cyan), anti-tubulin (red), Phalloidin (green), and DAPI (blue). (G) Confocal images of WT and cKO BMDCs cultured on non-coated or fibronectin-coated (10 μg/ml) glass slides and stained with anti-Paxillin (green) and DAPI (blue). White arrows indicate focal adhesion points. (H) Quantification (as mean fluorescence intensity, MFI) of CD11c and CD11b integrin receptors on the surface of WT and cKO BMDCs (n = 5 per genotype). Data are represented as mean ± SD. **p<0.01, ***p<0.001. (I) Immunoblot of CD-11c, FAM21 and actin in total cell lysates of WT and cKO BMDCs. (J) Trans-well migration assay (12-well) of WT and cKO BMDCs. Cells were seeded in the upper compartment of a trans-well plate and the chemokine CCL19 was added in the lower compartment (300 ng/ml) or non-treated (NT) and incubated for 3 h at 37 °C. Cells that migrated to the lower compartment were collected and counted (n = 3 per genotype). Data are represented as mean ± SD. *p<0.05.

FAM21 is known to attach to endosomes by binding to retromers and it recruits WASH to facilitate retrograde transport of vesicles through scission of cargo-enriched subdomains of early endosomes (Gomez & Billadeau, 2009; Harbour *et al*., 2012; Jia *et al*, 2010). We stained WT and FAM21-cKO BMDCs with anti-EEA1 (Figure 1D), measured ∼4000 endosomes in each cell population, we concluded that the early endosomes of cKO BMDCs were significantly enlarged relative to WT cells (Figure 1E). This outcome reflects a fission defect of endosomes in cKO cells, similar to our previous observation of FAM21 KO from HeLa cells (Hsiao *et al*., 2015).

Upon seeding WT and cKO BMDCs onto glass slides for imaging analyses, we noticed that most of the cKO cells remained unattached even after 16 h, whereas WT BMDCs were well attached, implying an adhesion defect of cKO cells (data not shown). Accordingly, we stained both cell types for cytoskeletal actin and α-tubulin, which revealed clear morphological differences between the WT and cKO cells (Figure 1F), with WT BMDCs being elongated and displaying cell polarity whereas cKO cells were rounded. Next, we seeded both cell types onto slides coated with fibronection and stained for focal adhesions using anti-paxillin antibody, which demonstrated that cKO cells hosted fewer focal adhesion points than WT (Figure 1G). We also noticed that surface expression of two integrin markers, CD11b and CD11c, was reduced in cKO cells relative to WT cells (Figure 1H), even though respective protein amounts in WT and cKO cell lysates were comparable (Figure 1I), implying an integrin receptor recycling defect similar to that described for WASH-KO cells (Duleh & Welch, 2012). Moreover, a cell migration assay revealed that cKO BMDCs moved faster than WT cells in the presence or absence of the chemokine CCL19 (Figure 1J). Taken together, these results indicate that FAM21-cKO cells exhibit altered cell morphology, diminished cell adhesion ability, and enhanced cell migration.

### FAM21 is important for the phagocytic activity of BMDCs

The immune-related functions of dendritic cells rely on their ability to take up foreign antigens, internally process them, and present them to other immune cells by displaying them on their surface. We tested if FAM21cKO affected the phagocytic function of BMDCs. WT and FAM21-cKO BMDCs were incubated with pHrodo-Red-labeled *E. coli* particles and then we analyzed (Figure 2A) and quantified (Figure 2B) the amounts of bacteria engulfed into endosomes by FACS at 45, 60 and 90 min after incubation, which showed that cKO cells displayed a reduced ability to phagocytose bacteria. Macropinocytic uptake of fluorescent dextran involves cellular engulfment of solutes from medium, and we also adopted this confocal microscopy approach at 30 and 60 min to show that dextran was internalized less efficiently in FAM21-cKO BMDCs compared to WT BMDCs (Figure 2C). Thus, we conclude that FAM21 is an important regulator of the endocytic activity of BMDCs.

**Figure 2:**
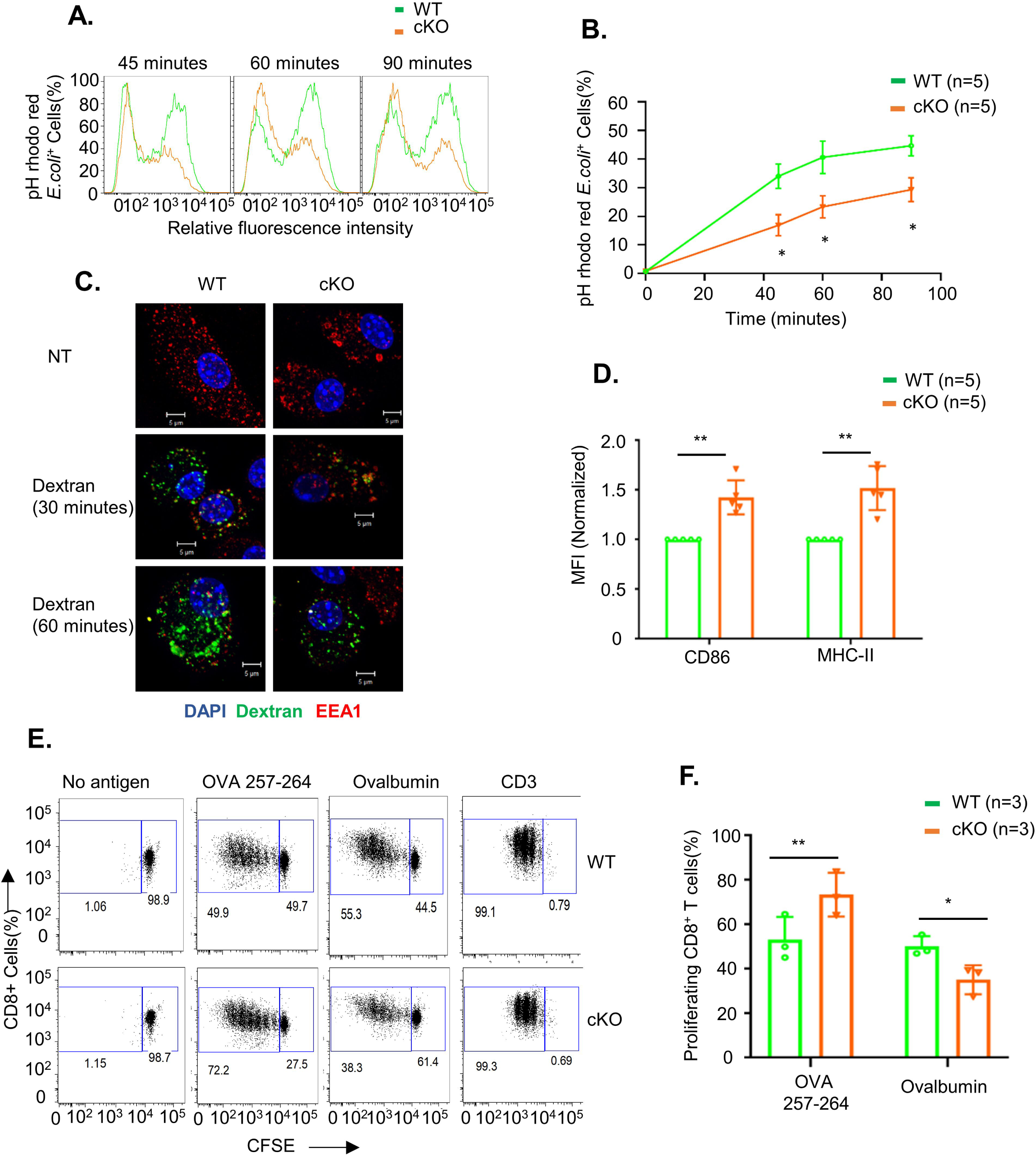
FAM21 is important for phagocytosis and the antigen processing function of BMDCs. (A) Fluorescence intensity of pHrodo Red *E.coli* taken up by WT and cKO BMDCs at 45, 60 and 90 min after adding bacteria to cells. (B) Percentage of BMDCs containing phagocytosed pHrodo Red *E.coli* from 0-90 min (n = 5 per genotype). Data are represented as mean ± SEM. *p<0.05. (C) Imaging analysis of dextran macropinocytosis in WT and cKO BMDCs. Cells were incubated with fluorescent dextran (green) (1 μg/ml) for 30 and 60 min at 37 °C, fixed, and then stained with anti-EEA1 (red) and DAPI (blue). NT; non-treatment control. (D) Quantification of MFI of co-stimulatory molecules CD86 and MHC class-II on the surface of WT and cKO BMDCs. Data are represented as mean ± SD. **p<0.01. (E) Quantification of the MFI of CD86 and MHC class-II (n = 5 per genotype) for WT and KO BMDCs. (F) Antigen presentation assay on WT and cKO BMDCs. CFSE-labeled CD8^+^ T cells isolated from OT-1 transgenic mice were co-cultured with WT and cKO BMDCs that had undergone prior pulsing with either ovalbumin protein or OVA 257-264 peptide. After 48 h of co-culture, CFSE dye dilution on CD8^+^ OT-1 cells was analysed as an index of T cell proliferation. CD3 served as a postitive control. (F) Quantification of CFSE-labeled CD8 ^+^ T cell proliferation, as described in (E) (n = 3 per genotype). Data are represented as mean ± SD. *p<0.05; **p<0.01.

We determined that the co-stimulatory molecules CD86 and MHC class-II were upregulated in cKO BMDCs relative to WT BMDCs (Figure 2D). Hence, we investigated if FAM21 cKO affects the antigen presentation function of BMDCs. To do so, we used two antigen types: ovalbumin that needs to be internalized, processed and loaded onto the co-stimulatory molecules of dendritic cells for presentation to T cells; and the small peptide OVA 257-264 that can be exogenously loaded onto cell surface co-stimulatory molecules for direct presentation to T cells. First, we pulsed BMDCs with ovalbumin or OVA 257-264, and then co-cultured them with CD8^+^ T cells isolated from OT-1 transgenic mice. CD8^+^ T cell proliferation was then monitored by FACS (Figure 2E) (Quah *et al*, 2007). We found that WT BMDCs co-cultured with ovalbumin or OVA 257-264 stimulated robust CD8^+^ T cell proliferation. In contrast, whereas FAM21-cKO BMDCs pulsed with OVA 257-264 stimulated CD8^+^ T cell proliferation better than WT BMDCs, they were less effective when pulsed with ovalbumin. Thus, cKO BMDCs displayed a reduced antigen processing ability, even though peptide antigen presentation on cell surfaces remained intact (Figure 2F). Consequently, FAM21 is important for the antigen processing function of BMDCs.

### Microarray analysis reveals downregulation of the TLR2/CLEC4E signaling pathway in FAM21-cKO BMDCs

A previous study of pancreatic cancer cells revealed that FAM21 knockdown affected NF- κB-mediated gene expression (Deng *et al*., 2015) in a WASH-independent manner. Hence, we investigated differential gene expression in WT and FAM21-cKO BMDCs by performing an unbiased microarray analysis of RNA samples. Microarray data was analyzed in GeneSpring (version 12.6.1), which identified 98 signficantly upregulated (FC >1.5) and 148 downregulated (FC <1.5) genes in FAM21-cKO BMDCs (Figure 3A&B). Next, we performed pathway enrichment analysis and identified upregulated (Figure EV3) and downregulated pathways (Figure 3C) and, notably, that cytokine-cytokine receptor interaction was the most downregulated pathway in the cKO BMDCs (Figure 3C). By means of STRING pathway analysis (https://string-db.org/), we discovered that several cellular transcripts in the TLR2/CLEC4E signaling pathway were downregulated in cKO BMDCs relative to WT (Figure 3D), such as those encoding CLEC4E, TLR2, CXCL2, IL-1β, and TNF-α, as validated by qRT-PCR (Figure 3E).

**Figure 3:**
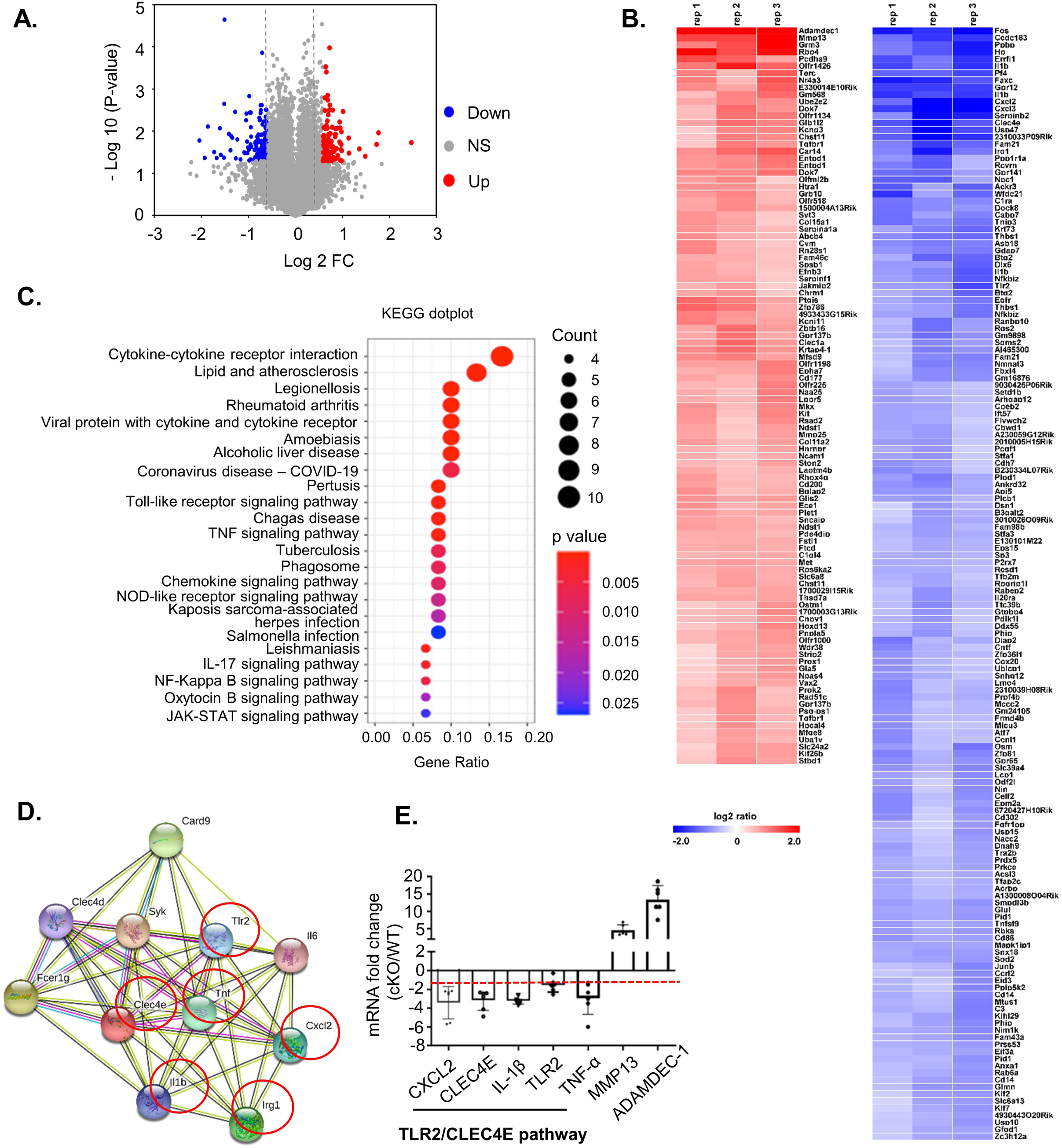
Microarray analysis reveals TLR2/CLEC4E pathway downregulation in cKO BMDCs. (A) Volcano plot depicting numbers of genes either 1.5-fold upregulated (red) or downregulated (blue) in cKO BMDCs relative to WT cells. All other genes are colored grey. (B) Heatmap of genes whose transcripts are either 1.5-fold upregulated (red) or downregulated (blue) in cKO BMDCs relative to WT BMDCs (n = 3 per genotype). (C) Gene enrichment analysis of the downregulated genes by KEGG dot plot. (D) STRING (Version 11.0) pathway analysis of CLEC4E-binding molecules. Red circles represent proteins that are downregulated >1.5-fold in cKO BMDCs relative to WT cells. (E) qRT-PCR of RNA samples from WT and cKO BMDCs (n = 6 per genotype). The red dashed line represents a 2-fold reduction level. Data are represented as mean ± SD. *p<0.05.

### FAM21 in BMDCs is important for TLR2/CLEC4E signaling pathway activation upon *C. albicans* infection

Next, we monitored cell surface expression levels of CLEC4E by flow cytometry (Figure 4A), with quantification demonstrating reduced levels of CLEC4E in FAM21-cKO BMDCs (Figure 4B). CLEC4E is a cell surface receptor expressed predominantly in myeloid cells such as monocytes, macrophages, and neutrophils {Matsumoto, 1999 #63;Ostrop, 2015 #65;Drouin, 2020 #64}. Together with TLR2, it is an important component of the pathogen- sensing cellular mechanism that detects infectious agents such as *C. albicans* and Mycobacteria, as well as other threats such as dead cell-derived proteins like SAP-130 {Wells, 2008 #55;Yamasaki, 2008 #56}. We infected WT and FAM21-cKO BMDCs with *C. albicans* and then measured phagocytosis of this pathogen at 15 and 30 min. Significantly, the cKO BMDCs phagocytosed much fewer *C. albicans* than WT BMDCs (Figure 4C&D). Moreover, phosphorylation of the *C. albicans*-induced downstream kinase SYK was attenuated in the cKO BMDCs relative to WT. We also stimulated the BMDCs with the ligand Zymosan, which is detected by dectin-1 and activates SYK phosphorylation. However, levels of SYK phosphorylation upon Zymosan treatment were similar between WT and FAM21 cKO BMDCs, suggesting CLEC4E-specific SYK downregulation in our cKO cells (Figure 4E). In addition, we tested a panel of innate immune stimuli—including HKMT and *C. albicans* as CLEC4E/TLR2-specific ligands, LPS as a TLR4-specific ligand, and Pam3SK as a TLR2-specific ligand—and then measured the expression of downstream cytokines {Ostrop, 2015 #65;Wells, 2008 #60;Clement, 2016 #66}. Our data in general support that the CLEC4E/TLR2 signaling pathway is considerably impaired in the cKO cells, whereas the LPS-induced TLR4 signaling pathway remains intact (Figure 4F&G). Together, these findings show that FAM21 protein is involved in the innate immune TLR2/CLEC4E signaling pathway of murine BMDCs, as summerized in Figure 4H.

**Figure 4:**
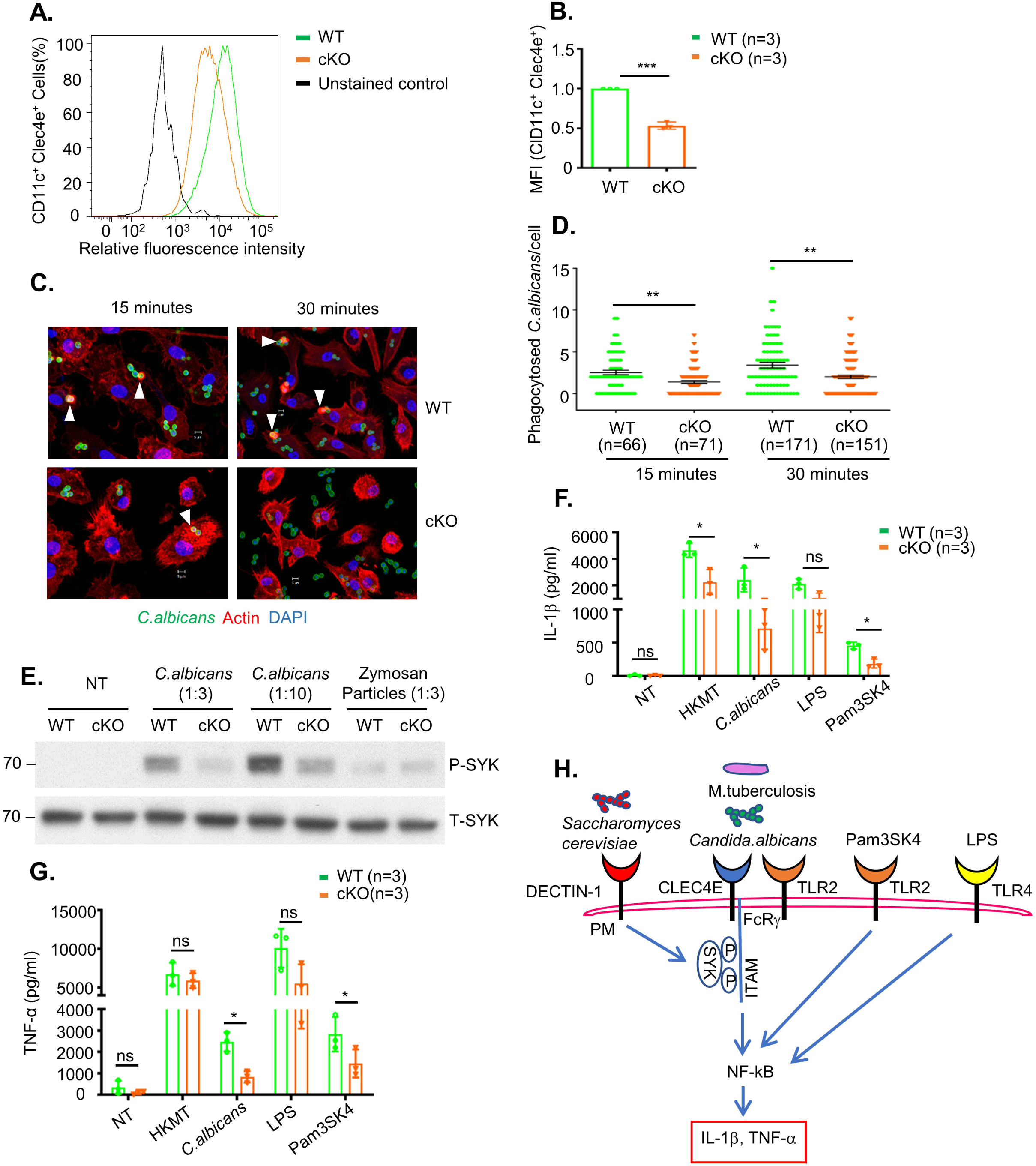
The TLR2/CLEC4E signaling pathway is downregulated in cKO BMDCs upon stimulation with specific ligands. (A) Surface expression of CLEC4E in CD11c^+^ BMDCs from WT and cKO mice, as determined by flow cytometry. (B) Quantification of CLEC4E MFI in WT and cKO BMDCs from (A) (n = 3 per genotype). (C) Representative images of phagocytosed *C. albicans* in WT and cKO BMDCs. CFSE-labeled *C. albicans* (green) was co-cultured with WT and FAM21-cKO BMDCs for 15 or 30 min, fixed with 4% paraformaldehyde, and then stained for Phalloidin (red) and DAPI (blue). Arrowheads point to phagocytosed *C. albicans.* (D) Quantification of phagocytosed *C. albicans* per cell for WT and cKO BMDCs from (C) (n, numbers of cells analyzed for each genotype for each timepoint). Data are represented as mean ± SEM. **p<0.01. (E) Immunoblot of phosphorylated SYK (P-SYK) and total SYK (T-SYK) in WT and cKO BMDCs at 30 min after infection with *C. albicans* and Zymosan particles. (F&G) ELISA of IL-1β (F) and TNF**-** α (G) levels in the supernatant of BMDCs at 16 h after being treated with HKMT, *C. albicans*, LPS and Pam3SK4 (n = 3 per genotype). (H) Graphical representation of signaling pathways affected in FAM21 cKO BMDCs compared to WT. Data are represented as mean ± SD. ns - not significant; *p<0.05.

### FAM21-cKO mice exhibit increased susceptibility to *C. albicans in vivo*

Next, we challenged WT and FAM21-cKO mice intraperitoneally with *C. albicans* and monitored body weight changes for 7-8 days. The cKO mice clearly lost more weight than WT mice between days 5-7 (Figure 5A), and 70% of the cKO mice had died by day 7 compared to only 20% of WT mice (Figure 5B). *C. albicans* titres in the kidneys of cKO mice were also higher than those in WT mice (Figure 5C). Moreover, H&E staining of kidneys at day 7 also revealed more *C. albicans* colonies in the cKO mice than in WT mice (Figure 5D), demonstrating that FAM21 functioning in CD11c^+^ dendritic cells is critical for *C. albicans* clearance *in vivo*.

**Figure 5:**
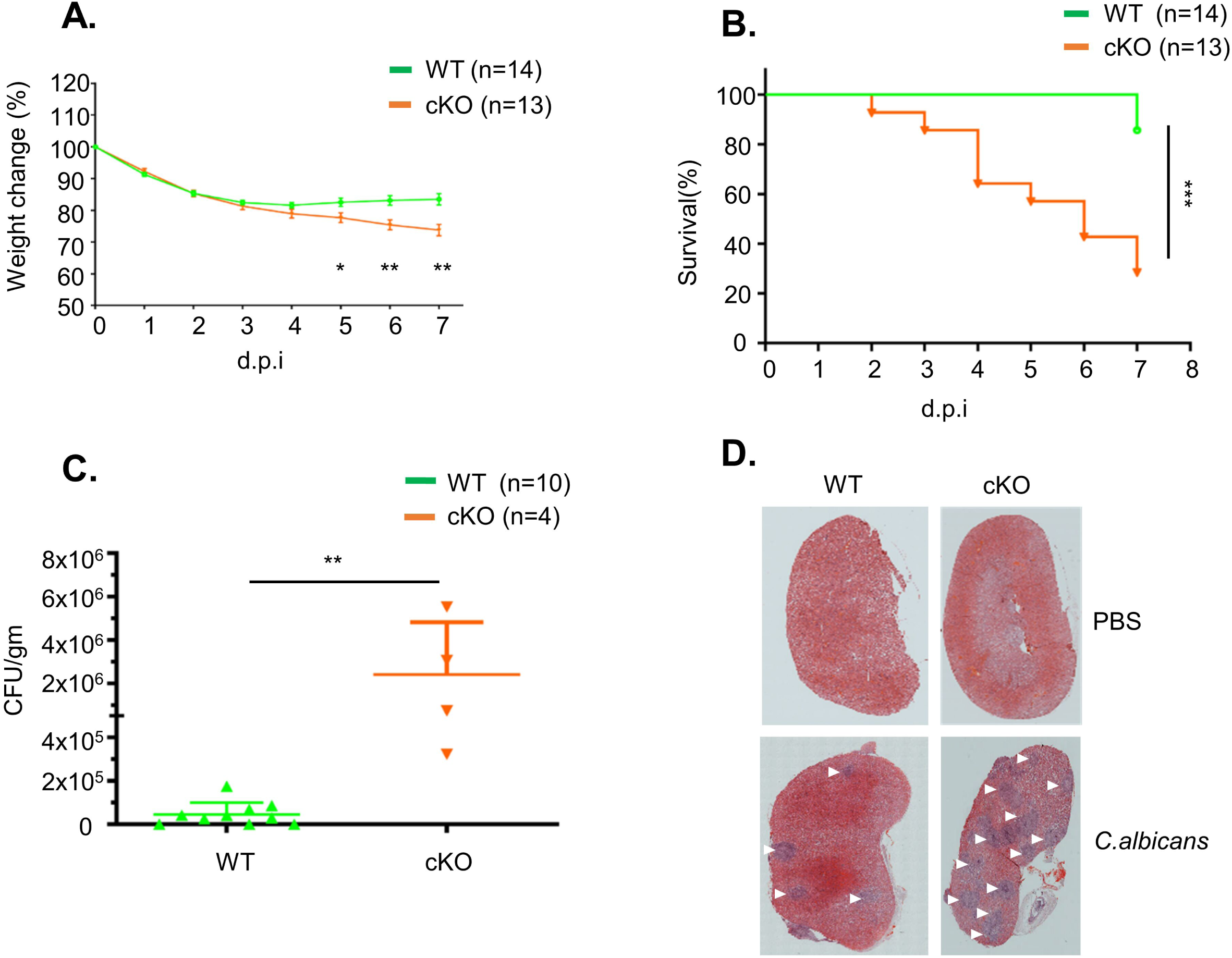
FAM21-cKO mice are susceptible to *C. albicans*. (A) Percentage of body weight change in WT and FAM21-cKO mice after intraperitoneal infection with 2.5x10^7^ *C. albicans* (n=14 for WT, n=13 for cKO). Data are represented as mean ± SEM. (B) Survival curves of the same mice described in (A). (C) Titers of *C. albicans* in the kidneys of infected mice at 7 d.p.i. (n=10 for WT, n=4 for cKO). Data are represented as mean ± SD. (D) Representative H&E staining of kidneys removed from infected WT and cKO mice at 7 d.p.i. White arrows mark the lesions caused by *C. albicans.* *p<0.05, **p<0.01, ***p<0.001.

### FAM21-cKO mice exhibit reduced innate and adaptive immune cell infiltration at infection sites

To understand if immune cell homeostasis is affected in FAM21-cKO mice upon *C. albicans* infection, we harvested immune cell populations from infection sites by means of intraperitoneal washes at days 1 and 3 post-infection (p.i.) for flow cytometry analyses (Figure EV4). At day 1 p.i., we detected reduced recruitment of dendritic cells (Figure 6A), macrophages (Figure 6B) and neutrophils (Figure 6C) in the FAM21-cKO mice relative to WT, but there were no major changes in their B and T cell populations (Figure 6 D-I). However, at day 3 d.p.i, recruitment of B and T cells, encompassing both CD4^+^ and CD8^+^ subsets, was significantly reduced in the cKO mice compared to WT (Figure 6 D-I), demonstrating that FAM21 cKO from CD11c^+^ dendritic cells of mice impacts multiple innate immune cell functions, as well as downstream B and T cell activation, upon *C. albicans* infection.

**Figure 6:**
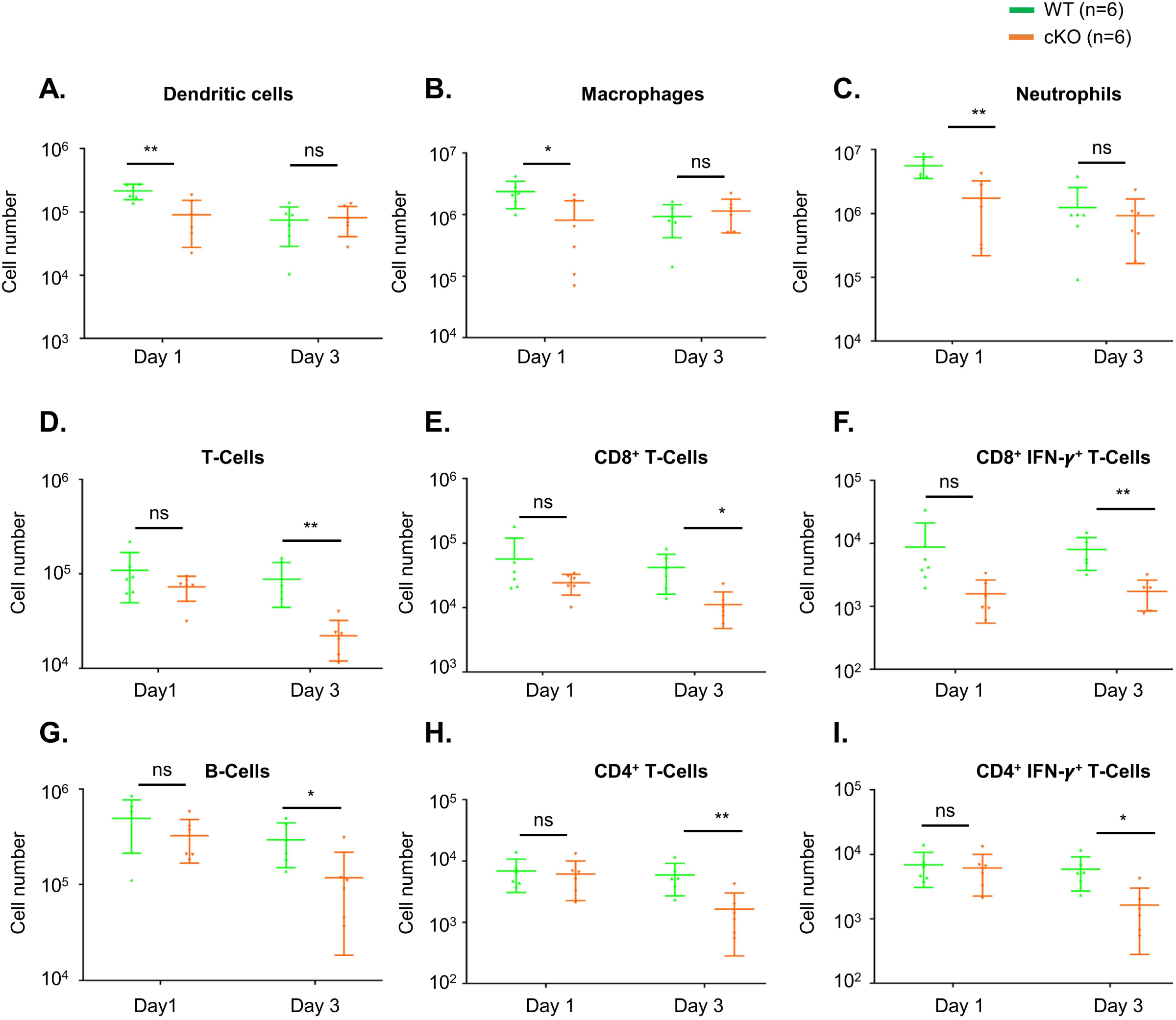
FAM21-cKO mice display defective innate and adaptive immune responses at the intraperitoneal site of *C. albicans* infection. Flow cytometry of intraperitoneal immune cell composition isolated from WT and cKO mice, as described in Figure EV4, at 1 and 3 d.p.i. with *C. albicans*. (A) Dendritic cells, (B) macrophages, (C) neutrophils, (D) total T cells, (E) CD8^+^ T cells, (F) CD8^+^IFNγ^+^ T cells, (G) B cells, (H) CD4^+^ T cells, and (I) CD4^+^IFNγ^+^ T cells (n=6 mice per genotype). Data are represented as mean ± SD. ns-not significant; *p<0.05; **p<0.01.

### Transfer of WT BMDCs *in vivo* partially protects FAM21-cKO mice against *C. albicans* infection

To demonstrate that the enhanced virulence of *C. albicans* in FAM21-cKO mice was indeed due to impaired CD11c^+^ dendritic cell function, we performed an adoptive transfer experiment whereby *in vitro*-cultured WT BMDCs were transferred intraperitoneally into either WT or FAM21-cKO mice 24 h prior to *C. albicans* infection. Following transfer, the WT and cKO mice were infected intraperitoneally with 2.5x10^7^ *C. albicans*, and then weight loss and survival were monitored. As shown in Figure 7A-D, the FAM21-cKO mice subjected to mock transfer (PBS injection) presented significant weight loss (Figure 7B) and lethality (Figure 7D), whereas the cKO mice pre-treated with WT BMDCs were well protected and all survived (Figure 7B&D). Control WT mice that received either PBS or WT- BMDCs presented minimal weight loss (Figure 7A) and all survived after infection (Figure 7C). Accordingly, we conclude that FAM21 is critical to how CD11c^+^ dendritic cells function to protect against *C. albicans* infection.

**Figure 7:**
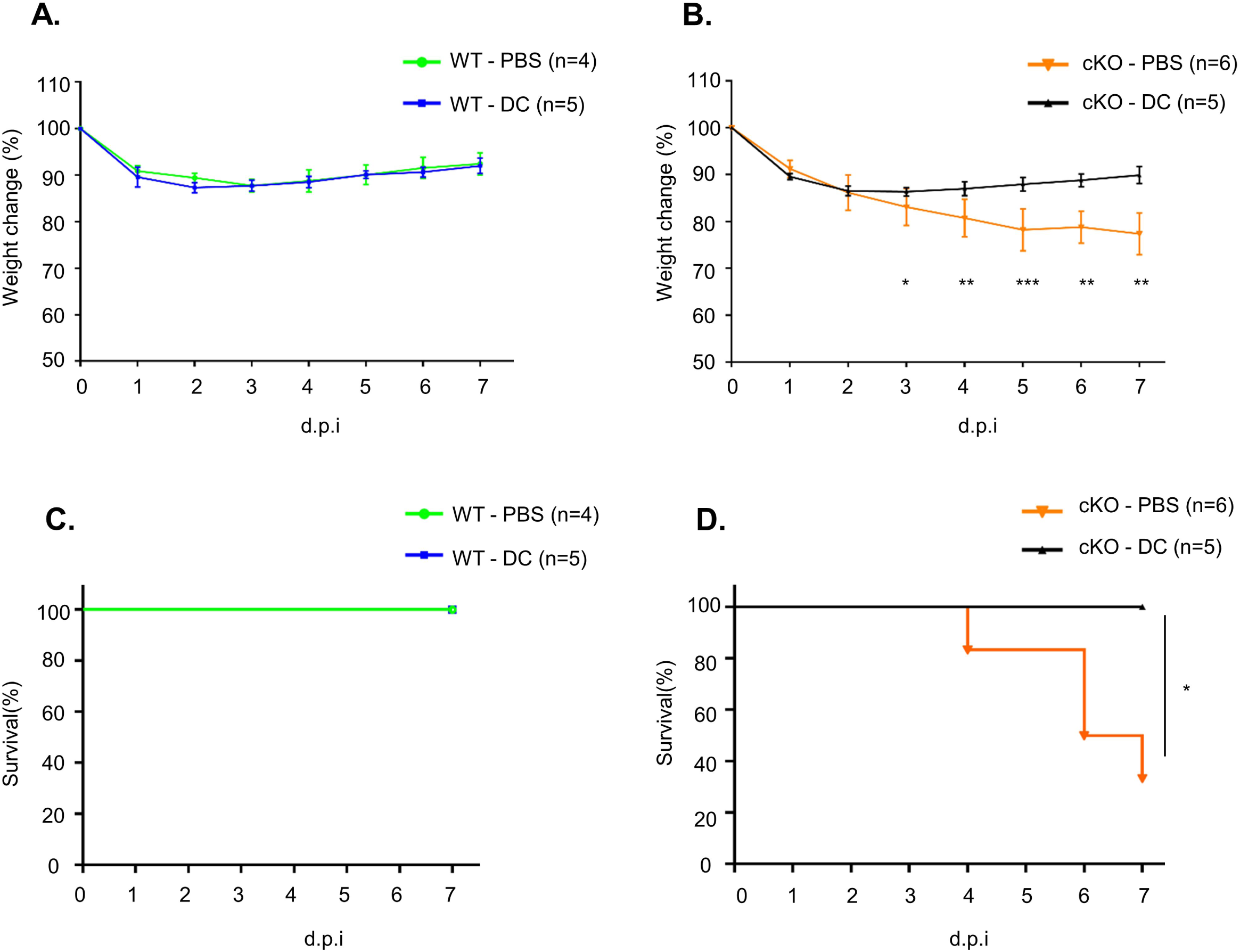
WT BMDC transfer rescues FAM21-cKO mice from the consequences of *C. albicans* infection. Weight change in (A) WT and (B) FAM21-cKO mice that were intraperitoneally administered with PBS or 5x10^5^ WT BMDCs (DC) prior to *C. albicans* infection. Data are represented as mean ± SD. (C&D) Survival curves at 7 d.p.i. of (C) WT mice described in (A), and (D) cKO mice described in (B) (n - represents numbers of mice in each group). * p<0.05, **p<0.01, ***p<0.001.

## Discussion

FAM21 is expressed in many mouse organs (Fig. EV1C) and our endeavors to generate constitutive KO mice were unsuccessful due to embryonic lethality (Figure EV1B). We observed that FAM21 is also expressed in BMDCs, which mediate important innate immune functions such as pathogen sensing, phagocytosis, and cytokine secretion, as well as antigen presentation to activate adaptive immune responses (Cabeza-Cabrerizo *et al*, 2021; Eisenbarth, 2019; Flannagan *et al*, 2012). Therefore, we generated FAM21-cKO mice with FAM21 gene deletion in the CD11C^+^ dendritic cell population. Previous studies have linked several components of the WASH complex to human diseases (Phillips-Krawczak *et al*., 2015; Song *et al*, 2018; Turk *et al*, 2017; Zavodszky *et al*, 2014), so our FAM21-cKO mice may serve as a useful animal model for further exploring human diseases and the future development of respective therapeutics.

Our *in vitro* characterization of FAM21-cKO BMDCs revealed their altered cell morphology and migratory capability. Those cells also displayed enlarged endosomes, consistent with previous data on FAM21-knockdown HeLa cells (Hsiao *et al*., 2015). Most importantly, our FAM21-cKO BMDCs exhibited a reduced ability for pathogen internalization and intracellular protein antigen processing, even though external peptide antigen presentation remained intact. Interestingly, our FAM21-cKO BMDCs also exhibited a reduced ability to activate the TLR2/CLEC4E signaling pathway *in vitro* and, FAM21-cKO mice challenged with *C. albicans* lost more body weight and displayed greater mortality when compared to WT mice. Our analyses of immune cell recruitment *in vivo* during the course of *C. albicans* infection of FAM21-cKO mice revealed fewer neutrophils and macrophages at the infection site at day 1 p.i., and fewer B and T cells at day 3 p.i.. Reconstituting FAM21-cKO mice with WT CD11c^+^ BMDCs rescued the cKO mice from death caused by *C. albicans* infection, supporting that FAM21 is important for TLR2/CLEC4E-mediated signaling in CD11c^+^ dendritic cells for host protection against *C. albicans*.

We found that some phenotypic differences of FAM21-cKO cells relative to WT could be linked to WASH complex functions. For example, WASH and ARP2/3 regulate early endosome fission, and WASH KO cells were found to possess enlarged early endosomes (Duleh & Welch, 2010), i.e., similar to our finding for FAM21-cKO cells. Moreover, integrins are known to mediate cell adhesion and phagocytic functions (Lukacsi *et al*, 2017; Torres-Gomez *et al*, 2020), and an accumulation of integrins in the macrophage lysosomes of *Drosophila* WASH mutants resulted in cell spreading defects (Nagel *et al*, 2017), again similar to our data herein on FAM21-cKO BMDCs. We noticed that levels of WASH protein, but not retromer subunits VPS35 and VPS26, were somewhat reduced in the FAM21-cKO BMDCs (Figure 1C), consistent with the above-described characterizations.

We also report some unique phenotypes of FAM21-cKO cells that differ from previous studies. (Graham *et al*, 2014) showed that WASH knockout from dendritic cells resulted in reduced surface levels of MHC class-II, and that the WASH-KO dendritic cells exhibited a diminished peptide antigen presentation capability (Graham *et al*., 2014). In contrast, our FAM21-cKO BMDCs displayed enhanced surface expression of MHC class-II antigen as well as peptide antigen presentation ability, relative to WT BMDCs. Thus, FAM21 and WASH may regulate MHC class-II recycling independently of each other or they function at different stages of the recycling pathway. Details of the underlying mechanism remain to be established.

A previous study reported that FAM21 in pancreatic cells may regulate NF-κB target gene transcription via chromatin binding in cell nuclei (Deng *et al*., 2015). Interestingly, CLEC4E has been reported as a target gene of the transcription factor C/EBPβ (Matsumoto *et al*., 1999). Hence, it is tempting to speculate that, besides its impact on NF-κB, impairment of the CLEC4E signaling pathway in FAM21-cKO BMDCs may be attributable to reduced chromatin binding of C/EBPβ to the promoter region of the *CLEC4E* gene. It will be interesting to elucidate if FAM21 regulates transcription factor interaction with chromatin in a WASH-independent manner. Overall, our study highlights the importance *in vivo* of FAM21 in host protection against pathogen invasion.

## Materials and methods

Our initial attempt to generate a constitutive FAM21 knockout (KO) mouse line failed because the respective KO mouse embryos exhibted abnormal development at embyronic day (E) 7.5, implying that FAM21 is essential for embryonesis (Figure EV1A&B). Accordingly, we altered strategy and generated FAM21-cKO mice using a BAC cloning system (RP23- 1904), obtained with the assistance of the Transgenic Core Facility of the Institute of Molecular Biology, Academia Sinica. We used the Red/ET BAC recombination system from Gene Bridges (Cat. Nos. K004 (loxP) and K005 (FRT)). Two loxP sites were inserted into introns 1 and 6 of FAM21 and a Frt-Neo-Frt selection cassette was then inserted after the loxP site in intron 1 according to the manufacturer’s instructions. We depict the targeting vector construct in Figure EV1C, in which 10 kilobasepairs (kbp) of the FAM21 gene sequence (Accession number: ENSMUSG00000024104) spanning exons 2 to 6 and encoding three in-frame methionine start codons were removed via Cre recombination. The resulting BAC construct was electroporated into C57BL6 embryonic stem (ES) cells (Open Biosystems). Three independent ES cell clones carrying the targeted allele were microinjected into albino C57BL/6J (Jackson Laboratory #00058) blastocysts to obtain FAM21^f-Neo/+^ germline-transmitted mice, which were then crossed with ACTB-FLPe mice to remove the Neo cassette. The resulting FAM21^f/+^ mice were then either mated with EIIa-Cre^+^ mice to obtain FAM21^+/-^ or were intercrossed with FAM21^f/+^ mice to obtain FAM21^f/f^. These latter homozygous FAM21^f/f^ floxed mice were crossed with CD11c-Cre^+^ mice to generate the FAM21^f/+^;CD11c-Cre^Tg/+^ mice, which were then crossed with FAM21^f/f^ mice to obtain the FAM21^f/f^;CD11c-Cre^Tg/+^line. Subsequently, FAM21^f/f^;CD11c-Cre^Tg/+^ mice were mated with FAM21^+/-^ to generate offspring of four different genotypes, i.e., FAM21^f/+^, FAM21^f/+^;CD11c-Cre^Tg/+^, FAM21^f/-^, and FAM21^f/-^;CD11c-Cre^Tg/+^ (Figure EV1E).

### Animal welfare

All animal protocols were approved by the Institutional Animal Care and Utilization Committee of Academia Sinica and were conducted in strict accordance with the guidelines on animal use and care of the Taiwan National Research Council’s Guide. Animals were sacrificed with carbon dioxide at the end-point of the experiment and all precautions were taken to minimize animal suffering throughout the study.

### BMDC cell culture

Murine BMDCs were cultured according to established protocols as described previously (Lutz *et al*, 1999), with minor modifications. In brief, bone marrow cells were isolated from the femur and tibia of 6-8-week-old WT and KO mice, counted, and seeded for 7 days at 2 x 10^6^ cells/10cm dish in bacterial culture plates in 10 ml conditioned medium [RPMI supplemented with 10% fetal bovine serum (FBS) (Hyclone), 1% penicillin-streptomycin (Gibco), 2mM L-Glutamine, 1mM Sodium pyruvate, 1% non-essential amino acids (Gibco), 20mM HEPES, 50μm 2-mercaptoethanol and 10% growth supernatant of GM-CSF transduced J558L cells (Qin *et al*, 1997). We replenished 10 ml of conditioned medium on day 3, and 50% of the medium was replaced with fresh medium on day 6. Non-adherent cells from the culture were collected on day 8 to analyse BMDC culture purity and for *in vitro* experiments. Generally, the cultured BMDCs comprised 80-85% CD11b^+^ CD11c^+^ cells, of which 10-15% were a CD86^+^ MHC class-II^+^ cell population according to flow cytometry analyses.

### Reagents and antibodies

Details of all reagents used in this study are presented in Table 1.

**Table 1:** Reagents and antibodies.

| Reagent name | Company | Catalog number |
| --- | --- | --- |
| Anti-WASH complex subunit FAM21C antibody | Merck | ABT79 |
| Anti-VPS26 antibody | Abcam | ab23892 |
| Anti-VPS35 antibody | Imgenex | IMG-3575 |
| Anti-WASH1 antibody | Sigma-Aldrich | SAB4200372 |
| Anti-Beta Actin antibody | BioVision | 3598-100 |
| Anti-EEA1 antibody | Santa Cruz | SC-6415 |
| Anti α-tubulin antibody | Amersham Life Science | N356 |
| Phalloidin-Tetramethylrhodamine B isothiocyanate | Sigma-Aldrich | P1951 |
| Anti-Phospho SYK (Tyr525/526) antibody | Cell Signaling | 2711 |
| Anti-SYK antibody | Cell Signaling | 2712 |
| Anti-CLEC4E/Mincle antibody | MBL | D292-3 |
| Anti-Paxillin antibody | BD Transduction Laboratories | 610051 |
| Cy5-Donkey anti-goat IgG (H+L) | Jackson Laboratories | 705-175-147 |
| Cy3-Donkey anti-rabbit IgG (H+L) | Jackson Laboratories | 711-166-152 |
| FITC-Goat anti-mouse IgG-Fab specific | Sigma-Aldrich | F8771 |
| Recombinant murine MIP-3β (CCL19) | PeproTech | 250-27B |
| pHrodo Red *E.coli* bioparticles | Molecular Probes | P35361 |
| Zymosan A *S.Cerevisiae* BioParticles, Texas<br>Red conjugate | Molecular Probes | Z2843 |
| Il-β mouse uncoated ELISA kit | Invitrogen | 88-7013 |
| Mouse TNF-α DuoSet ELISA | R&D Systems | DY410-05 |
| Fixable Viability dye eFluor 506 | eBioscience | 65-0866 |
| Heat-killed *Mycobacterium tuberculosis* | InvivoGen | tlrl-hkmt-1 |
| Lipopolysaccharides from *Escherichia coli*<br>0111:B4 | Sigma-Aldrich | L4391-1MG |
| Pam3CSK4 | InvivoGen | tlrl-pms |
| DAPI, Fluoropure grade | ThermoFisher<br>Scientific | D21490 |
| Albumin from chicken egg white | Sigma-Aldrich | A5503-10G |
| Dextran-Fluorescein | Invitrogen | D1820 |
| PE-anti-mouse CD11c antibody | BioLegend | 117308 |
| Pacific Blue anti-mouse I-A/I-E antibody | BioLegend | 107620 |
| Anti-mouse CD11b APC-eFlour 780 antibody | eBioscience | 47-0112-82 |
| CD86 (B7-2) Monoclonal antibody (GL1),<br>APC | eBioscience | 17-0862-82 |
| PE/Cyanine7 anti-mouse TCR $\beta$ chain antibody | BioLegend | 109222 |
| PerCP/Cyanine5.5 anti-mouse CD19 antibody | BioLegend | 115534 |
| Pacific Blue anti-mouse CD8α antibody | BioLegend | 100725 |
| PE/Cyanine7 anti-mouse CD4 antibody | BioLegend | 100422 |
| APC anti-mouse IFN-γ antibody | BioLegend | 505810 |
| PerCP/Cyanine5.5 anti-mouse NK-1.1 antibody | BioLegend | 108728 |
| PerCP/Cyanine5.5 anti-mouse CD3ε antibody | BioLegend | 100328 |
| APC anti-mouse F4/80 antibody | BioLegend | 123116 |
| FITC anti-mouse Ly-6C antibody | BioLegend | 128006 |
| PE/Cyanine7 anti-mouse Ly-6G antibody | BioLegend | 127618 |
| Alexa Fluor 488 anti-mouse IL-17A antibody | BioLegend | 506910 |
| ROR gamma (t) monoclonal antibody (B2D),<br>PE | eBioscience | 12-6918-82 |
| CD45.2 monoclonal antibody (104), PE-eFluor<br>610 | eBioscience | 61-0454-82 |

### Flow cytometry of BMDCs

Flow cytometry of BMDCs was performed as described in (Yin *et al*, 2015), with some modifications. In brief, 5x10^5^ BMDCs were preincubated for 20 min on ice with supernatant collected from hybridoma clone 2.4G2 cell culture (Unkeless, 1979) to block nonspecific binding of Fc gamma receptor, followed by staining for 20 min on ice with a fluorescent antibody cocktail containing anti-CD11c-PE (BioLegend), anti-CD11b-APC-eflour780 (eBiosciences), anti-CD86-APC (eBiosciences) anti-MHC class-II-PB (BioLegend). After washing three times with FACS buffer (PBS containing 2% FBS) and resuspension in FACS buffer, the BMDCs were analyzed by flow cytometry (BD LSR-II, BD Biosciences).

### Immunoblot analyses

Immunoblot analyses were performed as described previously (Kasani *et al*, 2017). In brief, 2x10^6^ BMDCs were collected, washed three times with PBS, and lysed in 100 µl sample buffer (4% SDS, 20% glycerol, 125 mM Tris pH 6.8, 0.25% bromophenol blue, 200 mM β- mercaptoethanol). Cell lysates were heated and separated by sodium dodecyl sulfate-polyacrylamide gel electrophoresis (SDS-PAGE), transferred to nitrocellulose membrane, and probed with antibodies recognizing FAM21 (Merck, 1:1000), WASH (Abcam, 1:1000), VPS29 (IMGENEX, 1:1000), VPS35 (Sigma, 1:1000), β-actin (BioVision, 1:2000) and CD11c-biotin (1:1000). To detect spleen tyrosine kinase (SYK) phosphorylation, 2x10^6^ BMDCs were infected with *C. albicans* at a ratio of 1:10 (cells:*C. albicans*) for 15 min at 37 °C and then washed three times with PBS. Cells were lysed using protein lysis buffer before loading equal amounts of protein (15 µg) per lane for SDS-PAGE. After transfer to nitrocellulose membrane, the membranes were blocked in 5% bovine serum albumin (BSA) and incubated overnight at 4 °C with primary anti-phospho-SYK (Cell Signaling Technologies, 1:1000) or anti-SYK (Cell Signaling Technologies, 1:1000) antibodies. The blots were then washed three times with PBST (PBS containing 0.1% Tween-20), incubated at room temperature for 1 h with HRP goat anti-rabbit IgG (Jackson, 1:20,000), and then developed using a Western Lightening Enhanced Chemiluminescence kit (PerkinElmer) according to the manufacturer’s protocol.

### Immunofluorescence imaging analyses

Immunofluorescence staining was performed as described previously (Izmailyan *et al*, 2012). In brief, 2x10^5^ BMDCs were seeded overnight on a glass coverslip in a 12-well plate with conditioned medium. The following day, the cells were fixed with 4% paraformaldehyde (PFA) at room temperature for 15 min, blocked with 1% BSA (Sigma) in PBS, and then stained with anti-EEA1 (Santa Cruz, 1:250), anti-FAM21 (Merck, 1:1000), anti-phalloidin- TRITC (Sigma, 1:1000) and anti-α-tubulin (Amersham, 1:1000) antibodies for 1 hour at room temperature in PBS containing 0.25% saponin and 1% BSA. The BMDCs were then stained with Cy5-Donkey anti-goat IgG (Jackson, 1:1000), Cy3-Donkey anti-rabbit IgG (H+L) (Jackson, 1:2000), FITC-Goat anti-mouse IgG-Fab specific (Sigma, 1:1000) secondary antibodies for 1 hour at room temperature. Nuclei were stained with DAPI for 5 min. Images were taken using a Zeiss LSM 780 confocal microscope with a 63x objective lens.

### Migration assay

The BMDC cell migration assay was performed as described previously (Scandella *et al*, 2004). In brief, 1x10^6^ BMDCs in 100 µl medium (RPMI + 1% FBS) were seeded in a well at the upper chamber of a 12-well trans-well plate (Costar, 3241), and 600 µl medium (RPMI + 1% FBS) containing the chemokine CCL19 (Peprotech) (300 ng/ml) was added to the lower chambers. The plates were incubated at 37 °C for 3 h, the number of cells that migrated through the membrane into the lower compartment were counted.

### Phagocytosis assay

Phagocytosis assay was performed by placing 2.5 x 10^5^ BMDCs in 100 μl on ice for 30 min and then transferring them to an Eppendorf tube. The medium was removed by centrifugation, and then the cells were incubated for 0-90 minutes at 37 °C with 100 µl of pHrodo Red *E.coli* (ThermoFisher, P35361). The BMDCs were washed three times with PBS and resuspended in FACS buffer, before being subjected to flow cytometry (BD LSR-II, BD Biosciences).

### Dextran uptake assay

WT and cKO BMDCs were incubated with Dextran-Fluorescein (Invitrogen, 10,000 MW) at a concentration of 1 μg/ml for 0-60 minutes at 37 °C, washed three times with PBS, and then fixed with 4% PFA. Next, the cells were stained with anti-EEA1 and DAPI, before being analyzed as described above in Immunofluorescence imaging analyses.

### Antigen presentation assay

Antigen presentation assay was performed as described previously (Wang *et al*, 2010) with some modifications. In brief, 1x10^5^ BMDCs were pulsed with ovalbumin protein (25 μg/ml) or with ovalbumin-derived OVA 257-264 peptide (SIINFEKL, 10 pm) for 16 h, washed three times with PBS, and then co-cultured with CD8^+^ T cells that were purified from spleen and lymph nodes of the OT-1 transgenic mouse and labeled with carboxyfluorescein succinimidyl ester (CFSE) at a concentration of 0.5 µM for 10 min (Pulle *et al*, 2006). Pulsed BMDC and CFSE-labeled CD8^+^ T cells co-cultured at a ratio of 1:5 for 48 h were sbubsequently washed with FACS buffer, stained with anti-CD8-APC antibodies, and analyzed by flow cytometry (BD LSR-II, BD Biosciences) to reveal CFSE dye dilution on CD8^+^ T cells.

### RNA extraction and labeling for microarray analysis

RNA was extracted from WT and cKO BMDC cultures using the TriZol (Invitrogen) method according to the manufacturer’s protocol. RNA quality assessment and labeling was performed as described previously (Kasani *et al*., 2017). In brief, 10 μg of RNA was reverse-transcribed with aminoallyl-modified dUTP using a Superscript Plus indirect cDNA labeling system (Invitrogen). The cDNA was then coupled with Alexa Fluor dye after purification using a Qiagen column. WT cDNA was labeled with Alexa Fluor 555, and cKO cDNAs were labelled with Alexa Fluor 647. These labeled cDNAs were hybridized to an Agilent SurePrint G3 Mouse GE 8 × 60K microarray (G4852A) according to the manufacturer’s protocol. The microarrays were scanned on an Agilent DNA microarray scanner (catalog number US9230696) using the two-color scan setting for 8 × 60K array slides. The scanned images were analyzed in Feature Extraction Software 10.5.1.1 (Agilent) using default parameters (protocol GE2_105_Dec08 and Grid 074809_D_F_20150624) to obtain background- and dye-Norm (Linear Lowess) signal intensities.

### Microarray and bioinformatic analyses

The raw microarray data (three biological replicates) were imported into GeneSpring software (Agilent, version 12.6.1). A total of 62,976 probes were then mapped to corresponding genes using the NCBI mm9 Mouse Genome Assembly (NCBI37; Jul2007). After removing probes associated with unnamed genes, non-coding genes or predicted genes, we were left with a total of 41,442 probes associated with 23,787 genes. Next, the probes with a signal intensity > 70 in at least two out of three replicates were selected to test if the log2 ratio of cKO/WT signals were significantly different from 0. Under the criteria of a Student *t*-test p-value < 0.05 and a fold-change > 1.5 or fold-change <1.5, we were left with 261 significant differentially-expressed probes representing 246 genes. Of these, 104 probes representing 98 genes were upregulated, whereas 157 probes representing 148 genes were downregulated in the FAM21-cKO mice. Next, we conducted Gene Ontology and KEGG pathway enrichment analysis on each set of genes using *clusterProfiler* (v 3.18.1) in the R package (Yu *et al*, 2012). Finally, the 148 downregulated genes were analyzed further for predicted and known protein-protein interactions using STRING (Version 11.0) (Szklarczyk *et al*, 2018).

### qRT-PCR analysis

RNA was isolated from WT and cKO BMDCs by the TriZol (Invitrogen) method. cDNAs were prepared from 1 µg RNA according to the manufacturer’s protocol. Quantitative real time-polymerase chain reaction (qRT-PCR) was performed on cDNAs using iQ SYBR green supermix (Bio-Rad) in a CFX-96 Touch system (Bio-Rad). Primer sequences are presented in Table 2.

**Table 2:**
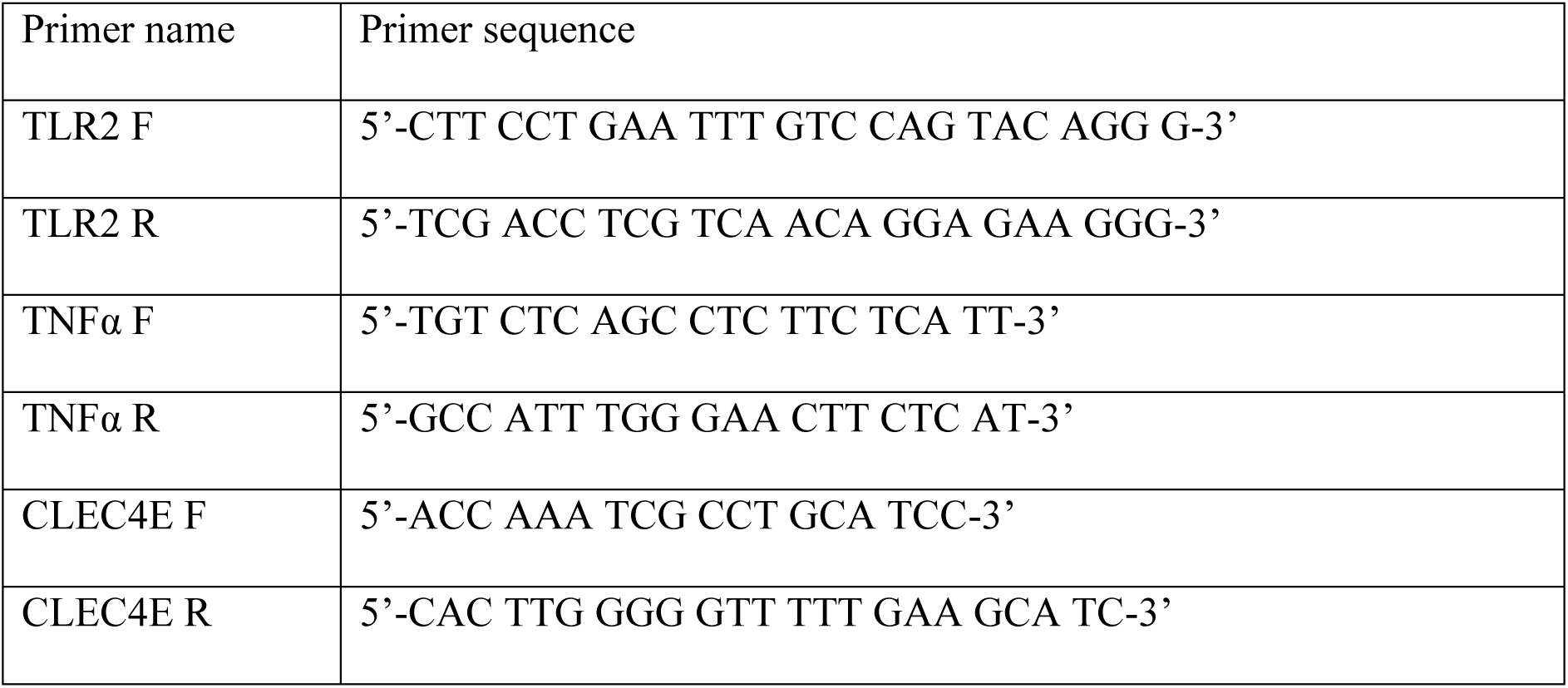
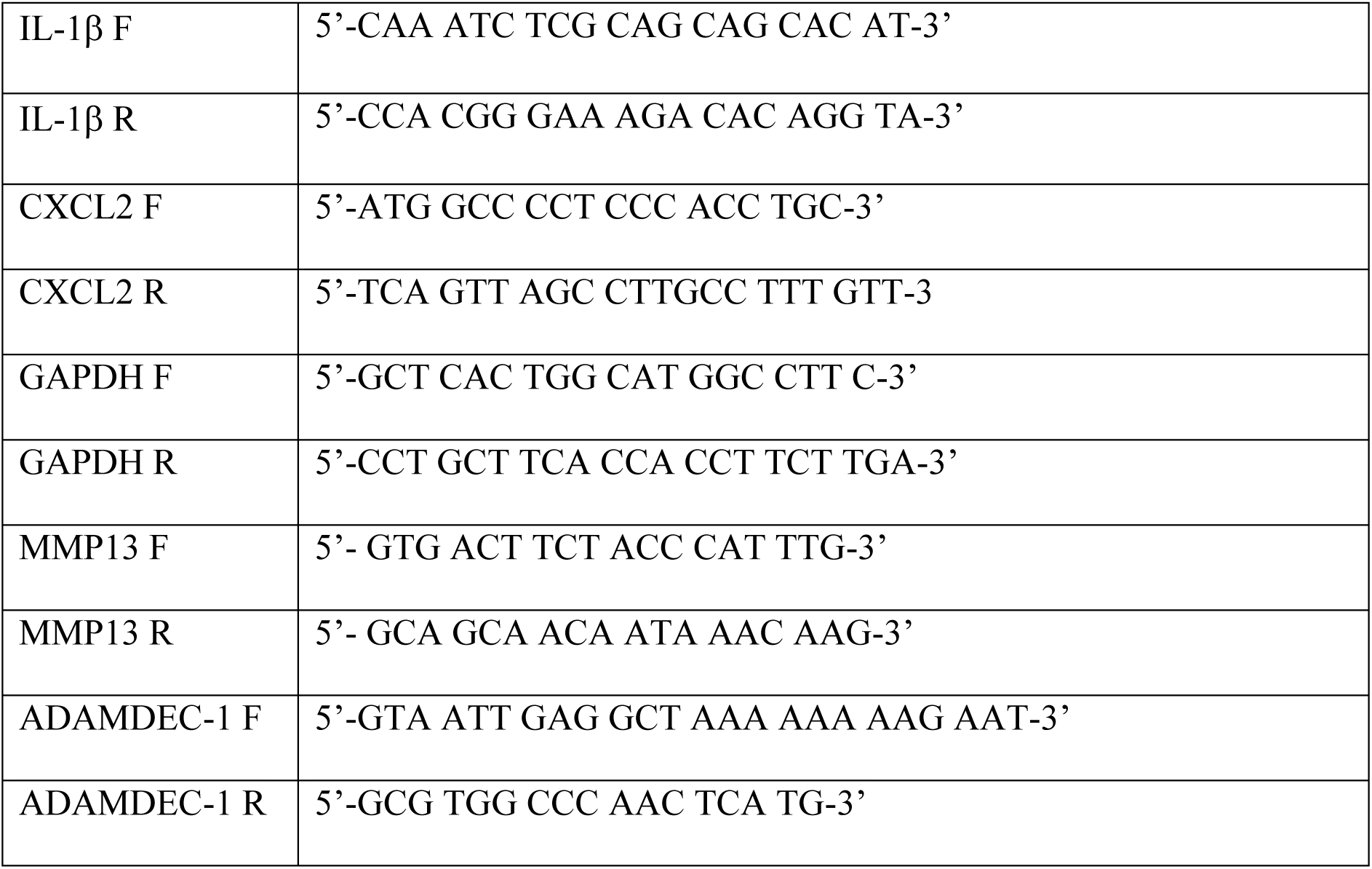
DNA primer sequences used in the study.

### Cytokine ELISA

WT and cKO BMDCs were seeded in a 96-well U-bottom plate (5x10^5^ cells/well) and stimulated for 16 h at 37 °C with each of the following ligands: lipopolysaccharide (LPS) (10 ng/ml), heat-killed *Mycobacterium tuberculosis* (HKMT, 5 µg/ml), PamCysSerLys (Pam3CSK4, 0.5 µg/ml) or *C. albicans* at a ratio 1:10 (cells:*C. albicans*) in 200 µl conditioned growth medium. Supernatants were collected and subjected to IL-1β (Invitrogen) and TNF-α (R&D) ELISA to detect those cytokines according to the manufacturers’ protocols.

### *C. albicans* uptake assay

We cultured 2x10^5^ BMDCs overnight on a glass slide in a 12-well plate and then incubated them for 15 or 30 min at 37 °C with CFSE-labeled *C. albicans* (ATCC 90028) (Dagher *et al*, 2018) at a ratio of 1:10 (cells:*C. albicans*). Glass slides were then washed three times with PBS and fixed with 4% PFA. The cells were stained for Phalloidin-TRITC (1:1000) and DAPI, and images were acquired using a Zeiss LSM 780 confocal microscope with a 63x objective lens.

### *In vivo* infection with *C. albicans* and intraperitoneal immune cell analysis

Male and female 8-9-week-old mice were intraperitonially (i.p) infected with 2.5x10^7^ *C. albicans* per mouse in 500 µl PBS as described previously (Ashizawa *et al*, 2019; Bergeron *et al*, 2017) and then monitored for disease progression by determining body weight loss. At 1, 3 and 6 days post-infection (d.p.i), mice were sacrificed and their intraperitoneal wash was harvested for immune cell population analysis by flow cytometry as described previously (Ray & Dittel, 2010). Mice were sacrificed when their body weight had been reduced by > 25% of their original body weight on the day of infection. The intraperitoneal wash cell suspension was treated with ammonium-chloride-potassium (ACK) lysis buffer to lyse the erythrocytes, followed by staining with a cocktail of fluorescent antibodies for myeloid and lymphoid cell populations (Table 1). Cells were analysed by flow cytometry (BD LSR-II, BD Biosciences).

### Histopathology

Mouse kidney histology was assessed as described previously (Kulkarni *et al*, 2021) with some modifications. The kidneys of WT and cKO mice were removed on 7 d.p.i and fixed in 4% paraformaldehyde for 2 days at 4 °C. The kidneys were then embedded, sectioned and stained with haematoxylin and eosin (H&E), followed by microscopic examination. Images were photographed using a Zeiss Axioimager-Z1 microscope with 20x objective lenses.

### BMDC transfer for *in vivo* complementation

We harvested 5x10^5^ WT BMDCs from culture on day 8 and transferred them intraperitoneally in 100 µl PBS into WT and cKO mice 24 h before *C. albicans* infection, as described previously (Burgess *et al*, 2014). The following day, mice were infected i.p. with 2.5x10^7^ CFU *C. albicans* per mouse. Mouse body weight was monitored for 7 days to observe the effect of cell complementation on *C. albicans* infection.

### Statistical analyses

Statistical analyses were performed using Student’s *t*-test in Graphpad prism software (version 9.0), and mice survival curves were generated using a Mantel-Cox test. p-values <0.05 were considered statistically significant: ns-not significant; *p<0.05; **p<0.01; ***p<0.001.

## Data availability

The data supporting the findings of this study are available within the manuscript and its supplementary materials. Microarray analysis data has been deposited to NCBI GEO with the GEO accession number: GSE196465.

## Acknowledgements

We thank Sue-Ping Lee of the Imaging Core Facility, Ya-Min Lin of the FACS Core Facility for the technical support at the Institute of Molecular Biology, Academia Sinica, Taipei, Taiwan. The work is supported by grants provided by Academia Sincia and Ministry of Science and Technology (110-2320-B-001 -015 -MY3).

## Author contributions

W.C conceived and supervised the project, W.C, R.K, S.K.K designed the experiments, R.K, S.K.K, S.Y.T, C.Y.T performed the experiments, R.K, S.K.K, K.H.Y, C.H.A.Y analysed the data, R.K, W.C wrote the manuscript, W.C provided grant support.

## Disclosure and competing intesrests

The authors declare no competing interests.

## The paper explained Problem

FAM21 protein, a component of WASH complex, is involved in endosomal fission of cargo containing vesicles in cells. However, despite its important role in cellular trafficking the *in vivo* function of FAM21 is unknown.

## Results

FAM21 is important for CD11c^+^ dendritic cell morphology and migration. Moreover, FAM21 is critical for phagocytosis and antigen processing function of these cells. Finally, FAM21 is required for the activation of TLR2/CLEC4E pathway and host protection against *C.albicans* infection.

## Impact

FAM21 is essential for CD11c^+^ dendritic cell function to activate host immune response against *C.albicans* infection.

**Figure EV1: Generation of FAM21-KO mice.** (A) Immunoblot analysis of FAM21^+/+^ and FAM21^-/-^ embryonic stem cells, as described in the text using anti-FAM21 and anti-β-actin antibodies. (B) WT^(+/+)^ and FAM21-KO^(-/-)^ embryos were isolated from uterine horns at E7.5, removed from yolk sacs, and photographed. (C) Northern blot showing ubiquitous expression of FAM21 mRNA in multiple organs of mice as well as in BMDCs. (D) A schematic representation of the targeting construct used to generate a floxed allele of the *FAM21* gene in mice via homologous recombination, followed by removal of the neomycin cassette. (E&F) A schematic representation of two mating strategies used to obtain FAM21-cKO mice, with specific deletion of FAM21 in CD11c^+^ cells for *in vitro* (E) and *in vivo* (F) experiments.

**Figure EV2: Three forms of FAM21 that may be derived from splicing variants.** Sketch map of FAM21 splice variants drawn from (http://asia.ensembl.org/Mus_musculus/Transcript/Summary?db=core;g=ENSMUSG00000024104;r=6:116184999-116239647;t=ENSMUST00000036759;tl=uDStZbFgkwFh7i2y-7980460). Comparision of the exon maps of three FAM21 splicing variants (not drawn to scale) and the respective proteins, with orange arrows representing FAM21 binding partners (not drawn to scale). (Freeman *et al*, 2014; Gomez & Billadeau, 2009; Harbour *et al*., 2012; Jia *et al*., 2010; Lee *et al*., 2016; McGough *et al*, 2014; Singla *et al*, 2019). * represents exons that were prematurely terminated compared to full length variant.

**Figure EV3: Enrichment analysis of the upregulated genes in FAM21-cKO BMDCs by KEGG dot plot.** KEGG dot plot enrichment analysis was performed on the gene sets that were upregulated (>1.5-fold) in FAM21 cKO BMDCs compared to WT BMDCs, showing the top signaling pathways that are significantly upregulated.

**Figure EV4: Characterization of the immune cells isolated from the intraperitoneal infection site.** Illustration of FACS analyses of intraperitoneal cells from C57BL/6 mice infected with *C. albicans* to identifiy (A) myeloid subsets including dendritic cells, macrophages and neutrophils, and (B) lymphoid subsets including B cells, T cells, CD4^+^ T cells, CD8^+^ T cells, CD4^+^IFNγ^+^ T cells and CD8^+^IFNγ^+^ T cells.

